# Structural heterogeneity of CTE Type I tau filaments with an interface-shifted polymorph

**DOI:** 10.64898/2026.09.10.750773

**Authors:** Ryohei Watanabe, Edward B. Lee

**Author notes:** Correspondence: Ryohei Watanabe, MD, PhD, Department of Psychiatry, University of Tsukuba Hospital, 2-1-1 Amakubo, Tsukuba, Ibaraki 305-8576, Japan.

## Abstract

Tau filament types can differ in inter-protofilament packing while sharing the same protofilament fold. Motivated by our recent findings in vacuolar tauopathy, we analyzed cryo-electron microscopy (cryo-EM) data from brain-derived chronic traumatic encephalopathy (CTE) tau filaments deposited in EMPIAR-10313. We resolved the canonical CTE Type I structure and a previously unreported CTE Type I-like structure with a shifted protofilament interface at 2.78 and 2.96 Å, respectively. Mapping particle-state assignments back onto the original micrographs showed that the CTE Type I and CTE Type I-like packing states can occupy locally contiguous regions within the same fibrils. These findings identify an interface-shifted CTE Type I-like structure and support its local coexistence with CTE Type I within individual fibrils.

## Introduction

Chronic traumatic encephalopathy (CTE) is a neurodegenerative tauopathy associated with repetitive head impacts and characterized neuropathologically by perivascular accumulation of phosphorylated tau, particularly at the depths of cortical sulci [1]. Cryo-electron microscopy (cryo-EM) has shown that CTE tau filaments comprise pairs of identical C-shaped protofilaments and that distinct filament types can arise from different inter-protofilament interfaces [2].

Our recent work on brain-derived tau filaments from vacuolar tauopathy (VT) examined several filament types, including VT Ia, which corresponds to the previously described CTE Type I core architecture [3,4]. Chemical repositioning of core-associated polyubiquitin with ubistatin B shifted the VT Ia protofilament interface, with the appearance of closely related interface-shifted structures [3]. Among the VT filament types analyzed in the same preparations, such interface rearrangement appeared to be restricted to VT Ia [3]. This selectivity raised the possibility that the paired-protofilament architecture shared by VT Ia and CTE Type I might likewise accommodate alternative inter-protofilament packing. We therefore asked whether such packing heterogeneity might also be present in brain-derived CTE Type I filaments.

To address this question, we examined cryo-EM data from brain-derived CTE tau filaments deposited in EMPIAR-10313 [2,5]. We identified, to our knowledge, a previously unreported CTE Type I-like structure with a shifted protofilament interface. Fibril-level back-projection further suggested that the CTE Type I and CTE Type I-like packing states can occupy locally contiguous regions within the same fibrils.

## Materials and Methods

### Public cryo-EM dataset and particle extraction

Deposited aligned micrographs and the coordinate-bearing particle file TypeI_particles.star were obtained from EMPIAR-10313 via the PDBj EMPIAR mirror [5]. Micrographs were imported into RELION 5.0.0 [6], and contrast transfer function (CTF) parameters were estimated with CTFFIND 4.1 [7]. Coordinates were retained only when the _rlnMicrographName entry in TypeI_particles.star matched a micrograph in the CTF-estimated list. Deposited coordinates were converted to RELION 5 per-micrograph coordinate STAR files while preserving the helical prior-related metadata; track-length values were converted from pixels to ångströms using the 1.055 Å/pixel sampling and stored as _rlnHelicalTrackLengthAngst, the standard RELION 5.0.0 helical metadata field for segment position along the helical track. Particles were extracted in 384-pixel boxes with a 300-Å tube diameter and two helical asymmetric units per segment. No additional coordinate-level selection was applied before extraction. The present study used only publicly available cryo-EM data and involved no new collection of human material; ethical approval and consent for the source study are described in Falcon et al. [2].

### Helical reconstruction and refinement

Reference-free 2D classification was performed in RELION 5.0.0 [6] with 100 classes and a regularization value of T = 1; particles in classes lacking recognizable filament features were discarded. Retained particles were subjected to 3D classification using an initial helical twist of −1.2°, a rise of 4.75 Å, a central Z fraction of 12%, and a VT IV reconstruction from our previous study, low-pass filtered to 40 Å, as the initial reference [3]. Selected 3D classes were iteratively processed by 3D auto-refinement, CTF refinement for beam tilt and higher-order aberrations, anisotropic magnification and defocus, and post-processing. Final 3D auto-refinements were performed with C1 symmetry. Overall resolutions were estimated from gold-standard Fourier shell correlation (FSC) between independently refined half-maps at FSC = 0.143. Local-resolution maps were calculated from the final half-maps using the corresponding solvent masks used for post-processing.

### Atomic model building, refinement and interface analysis

Initial atomic models were generated with ModelAngelo 1.0.18 from the final post-processed maps using the tau sequence (UniProt P10636-8) [8]. Models were manually adjusted to the corresponding maps in UCSF ChimeraX 1.10 [9]. Seven adjacent cross-β rungs were retained and refined against the corresponding post-processed maps by real-space refinement in PHENIX 1.20.1 [10]. Model–map FSC and half-map cross-validation were assessed in PHENIX; for cross-validation, models refined against half-map 1 were evaluated against the independent half-map 2 without further coordinate refinement. Inter-protofilament hydrogen-bonding contacts were identified in the refined seven-rung models using UCSF ChimeraX with identical relaxed geometric criteria for both structures (0.4 Å distance and 20° angle tolerances). Counts per cross-β rung were derived from the periodic contact pattern. Buried inter-protofilament solvent-accessible surface areas were calculated from the same seven-rung models using the standard buried-area function in ChimeraX with a 1.4-Å solvent probe. Model-refinement and validation statistics are summarized in Table S1.

### Fibril-state analysis by back-projection onto original micrographs

The fibril-state analysis used relion_fibril_state_analysis.py v1.0.0 on particle-level state assignments from the four-class RELION 5.0.0 3D classification. The three classes corresponding to CTE Type I were pooled as the CTE Type I state, whereas the remaining class was assigned as CTE Type I-like. Particle-level state assignments were mapped back onto the original aligned micrographs using the corresponding particle coordinates. For this analysis, a fibril was operationally defined by the unique combination of micrograph name and RELION helical-tube identifier, and particles were ordered by _rlnHelicalTrackLengthAngst. The nominal segment step was 14.1 Å; adjacent particles were considered contiguous at 14.1 ± 0.2 Å. For each fibril, the expected number of segment positions was calculated as round[(maximum track length − minimum track length)/14.1] + 1, and coverage was the number of observed particles divided by the expected number of positions. Only fibrils in which at least 95% of the expected segment positions were occupied by particles assigned to either CTE Type I or CTE Type I-like were included in the composition analysis. Fibrils were classified as pure CTE Type I, pure CTE Type I-like, or mixed according to the states of observed particles within the sampled track. Among mixed fibrils, a locally supported transition was defined as an adjacent state change between CTE Type I and CTE Type I-like with nominal spacing and at least five consecutively spaced same-state particles immediately flanking each side. Representative aligned-micrograph fields were selected with extract_candidate_micrograph_crops.py v1.0.0. Particle-state assignments and locally supported transition sites were projected onto the corresponding aligned micrographs with export_final_overlay_panels.py v1.0.0. Map/model panels were rendered in UCSF ChimeraX 1.10 [9].

## Results

The locally downloaded dataset comprised 5,303 aligned micrographs at 1.055 Å/pixel and 203,154 deposited CTE Type I segment coordinates. Matching the deposited coordinates to the CTF-estimated micrographs retained 198,298 segments from 1,898 micrographs (Fig. 1a). Following 2D classification, 192,484 particles were retained for 3D classification. 3D classification into four classes yielded three CTE Type I classes (150,228 particles in total) and one CTE Type I-like class (42,256 particles; 22.0%). The best-resolved CTE Type I class contained 33,415 particles. Final refinement of the CTE Type I and CTE Type I-like classes with C1 symmetry yielded a helical twist of 179.4° and a rise of 2.37 Å for both structures, with overall resolutions of 2.78 Å and 2.96 Å, respectively (Fig. 1b-d; Table S1).

**Fig. 1.**
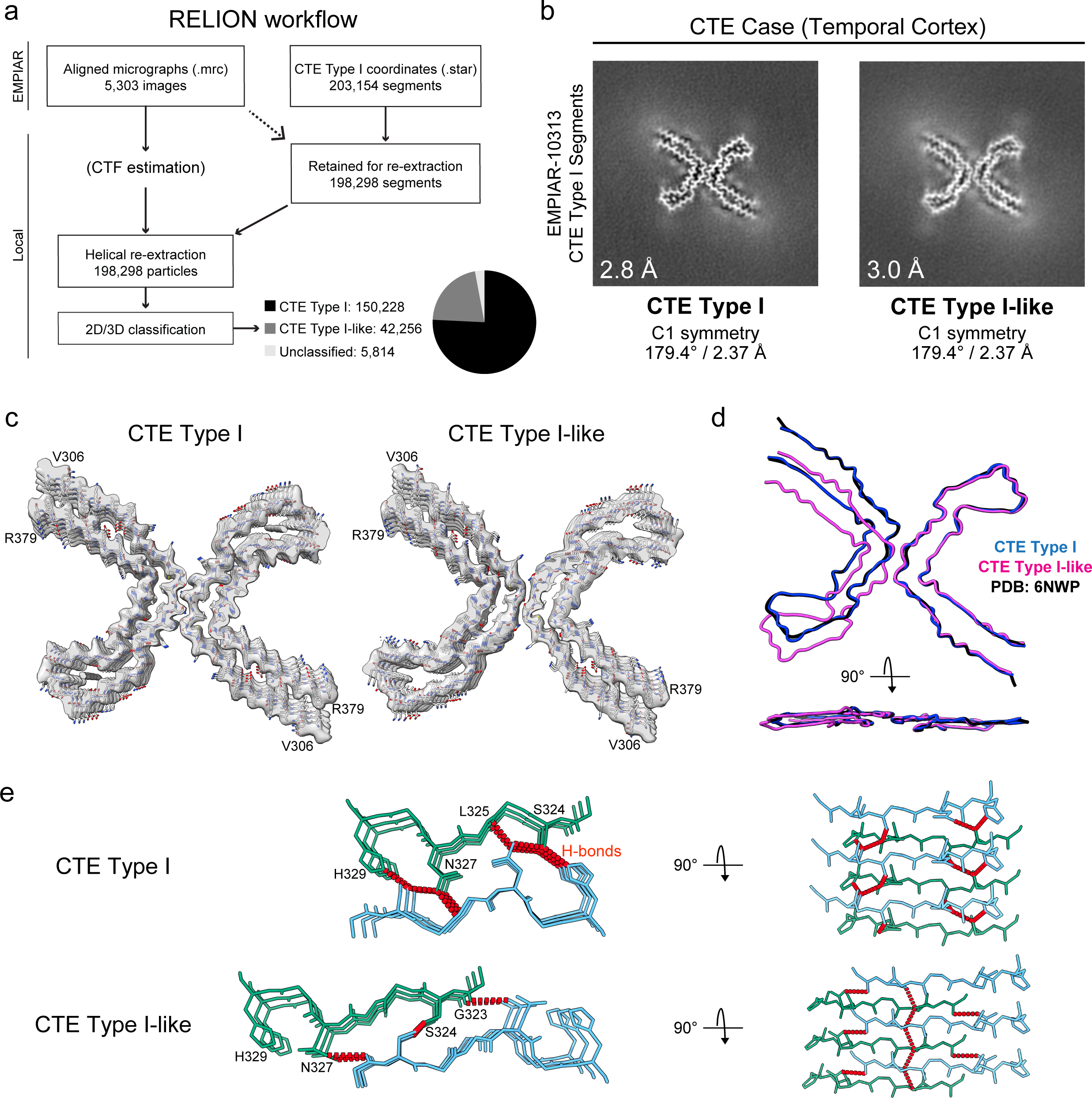
Structural identification and interface comparison of CTE Type I and CTE Type I-like tau filaments. (a) RELION workflow for analysis of EMPIAR-10313 [5]. Locally obtained aligned micrographs and deposited CTE Type I segment coordinates were used for helical re-extraction and 2D/3D classification. Of 203,154 deposited coordinates, 198,298 segments were retained for extraction. Following 2D classification, 192,484 particles were retained for 3D classification, yielding 150,228 CTE Type I and 42,256 CTE Type I-like particles. (b) Averaged XY cross-sections of the final reconstructed maps for CTE Type I and CTE Type I-like. Final overall resolutions were 2.78 Å and 2.96 Å, respectively. Both reconstructions were refined with C1 symmetry, with a helical twist of 179.4° and a rise of 2.37 Å. (c) Final reconstructed maps with the corresponding fitted atomic models. Both models comprise residues V306–R379 in each protofilament. (d) Overlay of the reconstructed CTE Type I (blue), CTE Type I-like (magenta), and published CTE Type I (PDB 6NWP; black) models in axial and 90°-rotated side views [2]. The protofilaments retain the same overall fold, whereas their relative packing is shifted in CTE Type I-like. (e) Close-up comparison of the inter-protofilament interfaces. The same G323–H329 region is shown for both structures, with backbone atoms displayed throughout and the side chains of S324, N327, and H329 shown in both structures for direct comparison. Red dashed lines indicate geometrically defined inter-protofilament hydrogen-bonding contacts. CTE Type I contains S324–H329, N327–S324, and N327–L325 contacts, whereas the shifted CTE Type I-like interface contains S324–S324 and N327–G323 contacts. Corresponding 90°-rotated views show three adjacent cross-β rungs and the axial organization of the hydrogen-bonding networks.

Refinement of the CTE Type I class produced a structure closely matching the published CTE Type I model (PDB 6NWP), whereas the CTE Type I-like class yielded a previously unreported brain-derived CTE filament structure [2,11]. Both refined models comprised residues V306– R379 in each protofilament and retained the same overall protofilament fold. In CTE Type I, the inter-protofilament interface was centered on residues G323–H329 and contained six inter-protofilament hydrogen-bonding contacts per cross-β rung, comprising two each of S324–H329, N327–S324, and N327–L325 contacts. In CTE Type I-like, relative displacement of the protofilaments shifted the principal interface toward residues G323–N327 and reorganized the network to four hydrogen-bonding contacts per cross-β rung, comprising two S324–S324 and two N327–G323 contacts, a 33% reduction. The matched seven-rung models also showed buried inter-protofilament solvent-accessible surface areas of 1,108 Å² for CTE Type I and 874 Å² for CTE Type I-like, corresponding to a 21% smaller buried interface in CTE Type I-like. Thus, the principal structural difference between the two structures lies in the relative protofilament packing and interaction pattern at the interface rather than in the underlying protofilament fold (Fig. 1c–e).

Model resolutions were 3.0 Å for CTE Type I and 3.2 Å for CTE Type I-like (model–map FSC = 0.5). MolProbity scores were 2.21 and 2.20, respectively, with no Ramachandran outliers (Table S1). Local-resolution maps showed comparable resolution at the inter-protofilament interfaces, and half-map and model–map FSC analyses supported both reconstructions (Fig. S1). The interface differences are therefore unlikely to result solely from the modestly lower map quality of CTE Type I-like.

Back-projection of particle-state assignments was then used to determine how the two packing states were distributed along individual fibrils (Fig. 2a). Among 2,337 high-coverage fibrils derived from the deposited CTE Type I segment set, with at least 95% coverage of the expected 14.1-Å-spaced segment positions, 1,006 (43.0%) were pure CTE Type I, 104 (4.5%) were pure CTE Type I-like, and 1,227 (52.5%) contained particles assigned to both states (Fig. 2b).

**Fig. 2.**
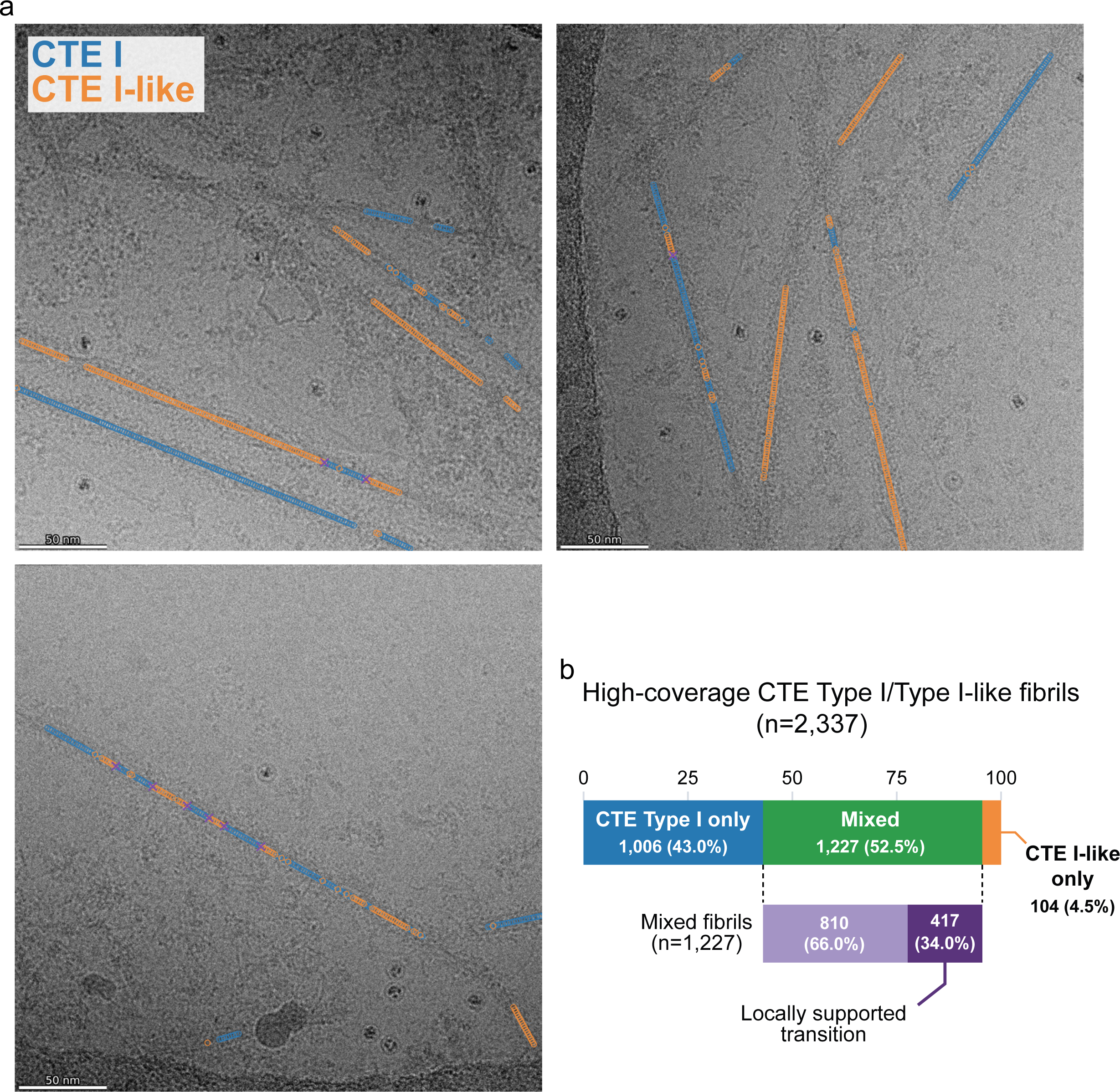
Fibril-level distribution of CTE Type I and CTE Type I-like packing states. (a) Representative aligned cryo-EM micrographs showing fibril-level back-projection of particle-state assignments. CTE Type I and CTE Type I-like assignments are shown in blue and orange, respectively. Purple marks denote locally supported transition sites. Scale bars, 50 nm. (b) Distribution of packing states among high-coverage fibrils, defined as fibrils with CTE Type I or CTE Type I-like assignments at ≥95% of the expected 14.1-Å-spaced segment positions. These fibrils were derived from the deposited CTE Type I segment set. Of 2,337 high-coverage fibrils, 1,006 (43.0%) contained only CTE Type I assignments, 104 (4.5%) contained only CTE Type I-like assignments, and 1,227 (52.5%) contained both states. Among the 1,227 mixed fibrils, 417 (34.0%) contained at least one locally supported transition, defined as an adjacent CTE Type I–CTE Type I-like state change at 14.1 ± 0.2 Å with at least five consecutively spaced same-state particles immediately flanking each side of the boundary; the remaining 810 (66.0%) did not meet this criterion.

To more stringently assess local continuity, an adjacent state change at the nominal segment spacing was required to be flanked by at least five consecutively spaced same-state particles on each side. Of the 1,227 mixed fibrils, 417 (34.0%; 17.8% of the high-coverage cohort) met this criterion (Fig. 2a,b), providing more stringent support that CTE Type I and CTE Type I-like packing states occur in locally contiguous regions within individual fibrils.

## Discussion

Our findings identify CTE Type I-like as a filament type closely related to CTE Type I in brain-derived CTE tau. The individual protofilaments retain the same overall fold, but their packing differs from that of CTE Type I. The three previously reported CTE filament types likewise share a common protofilament fold but differ in their inter-protofilament packing, with each reported as an axially repetitive structure with a single packing arrangement [2,11]. In our fibril-level analysis, back-projection supported local coexistence of the CTE Type I and CTE Type I-like structures as distinct packing states within individual tau fibrils. Taken together, these findings add to the known structural diversity of brain-derived tau filaments in CTE and support local variation in inter-protofilament packing along individual fibrils.

In cryo-EM studies of tau filaments from human neurodegenerative disease brains, structural polymorphism has generally been described in terms of distinct filament populations, even when multiple filament types coexist within the same disease or specimen [12,13]. Related fibril-level heterogeneity has been examined for brain-derived α-synuclein filaments from multiple system atrophy, where a minority of filaments showed mixed Type I_2_/II_2_ assignments. However, the authors favored classification stochasticity over true mixed filaments because the Type I_2_ and II_2_ protofilament folds were largely dissimilar and modeled adjacent Type I_2_/II_2_ rungs produced severe steric clashes [14]. By contrast, CTE Type I and CTE Type I-like retain the same overall protofilament fold and differ principally at the protofilament interface, making local changes in packing structurally more plausible. The presence of fibrils containing only CTE Type I-like assignments further indicates that this packing state is not confined to transition regions within mixed fibrils, compatible with maintenance of this packing state during fibril elongation. For mixed fibrils, we further required putative transitions to be flanked on each side by at least five consecutively spaced same-state particles, rather than relying on mixed assignments alone. Adjacent extracted segments overlap extensively, so consecutive state assignments cannot be treated as independent observations and the precise transition boundary cannot be resolved at atomic scale. Even so, the continuity of same-state assignments on both sides of putative transitions supports local variation in inter-protofilament packing within individual fibrils.

What determines this packing heterogeneity remains unclear. The CTE Type I interface, also observed in VT Ia in our previous work, lacks the specific electrostatic interactions between oppositely charged residues present in some other tau filament interfaces [2,3]. Because such electrostatic interactions can constrain the relative geometry of interacting residues [15], their absence may permit a broader range of inter-protofilament packing geometries. CTE Type I-like retains the same protofilament fold but forms a reorganized interface with fewer hydrogen-bonding contacts and a smaller buried surface area. A related rearrangement of the corresponding VT Ia interface was observed following chemical repositioning of core-associated polyubiquitin [3]. Time-resolved cryo-EM of recombinant tau revealed closely related intermediates with small differences in protofilament packing, suggesting relative protofilament sliding, potentially at filament ends or defects [16]. Such lateral rearrangement is also compatible with the mechanical anisotropy of amyloid fibrils, in which longitudinal hydrogen-bonding interactions are much stiffer than transverse interactions [17]. These observations suggest that the CTE Type I interface may permit more than one packing arrangement.

Fibril-level analyses of additional CTE cases and other diseases with CTE-fold tau filaments, including subacute sclerosing panencephalitis and amyotrophic lateral sclerosis/parkinsonism-dementia complex, could establish how broadly local packing heterogeneity occurs [11,18]. Recent cryo-electron tomography of postmortem Alzheimer’s disease brain showed that tau filament heterogeneity can be spatially organized within intact tissue [19]. Applying such in situ approaches to tau pathology in CTE may reveal whether the packing variation identified here is maintained in native tissue and how it relates to the surrounding cellular environment. More broadly, whether within-fibril structural heterogeneity is common among brain-derived tau filaments remains an open question; if so, it would represent an additional level of tau polymorphism beyond the discrete filament types typically resolved by helical reconstruction.

## Acknowledgements

We thank the EMPIAR team for maintaining a public archive that enabled this study. This study was supported by grants from the NIH (R01AG065341, P30AG072979, and P01AG066597 to EBL), the DeCrane Family Fund for PPA Research, and gifts from the Shanahan Family Foundation and the Barrist Family Foundation. The funders had no role in study design; data collection, analysis, or interpretation; preparation of the manuscript; or the decision to submit the article for publication. The authors used ChatGPT (OpenAI) to assist with manuscript reformatting and language editing. The authors reviewed and edited the generated output and take full responsibility for the content of the publication.

## Conflicts of interest

The authors declare no conflicts of interest.

## Author contributions

RW and EBL conceived the study. RW performed the cryo-EM data analysis and drafted the manuscript. RW and EBL reviewed and approved the final version.

## Data availability

The data that support the findings of this study were derived from the publicly available Electron Microscopy Public Image Archive dataset EMPIAR: EMPIAR-10313 (https://doi.org/10.6019/EMPIAR-10313). The cryo-EM maps generated in this study have been deposited in the Electron Microscopy Data Bank under EMDB: EMD-78521 (CTE Type I) and EMDB: EMD-78522 (CTE Type I-like). The refined atomic models have been deposited in the Protein Data Bank under PDB: 37VX (CTE Type I) and PDB: 37VY (CTE Type I-like).

## Code availability

The custom Python scripts and paper-specific configuration files used for the fibril-state analysis and figure generation have been deposited in Zenodo under the reserved DOI 10.5281/zenodo.21856527.

## Abbreviations

CTE: chronic traumatic encephalopathy
CTF: contrast transfer function
cryo-EM: cryo-electron microscopy
FSC: Fourier shell correlation
VT: vacuolar tauopathy

## Notes

https://doi.org/10.6019/EMPIAR-10313

### Competing Interest Statement

The authors have declared no competing interest.

### Summary of Updates

Figure 1b was corrected by replacing the two map panels with the intended images. No other changes were made.

