## Supplementary Information for "Structural heterogeneity of CTE Type I tau filaments with an interface-shifted polymorph"

**This file contains:**

Fig. S1

Table S1

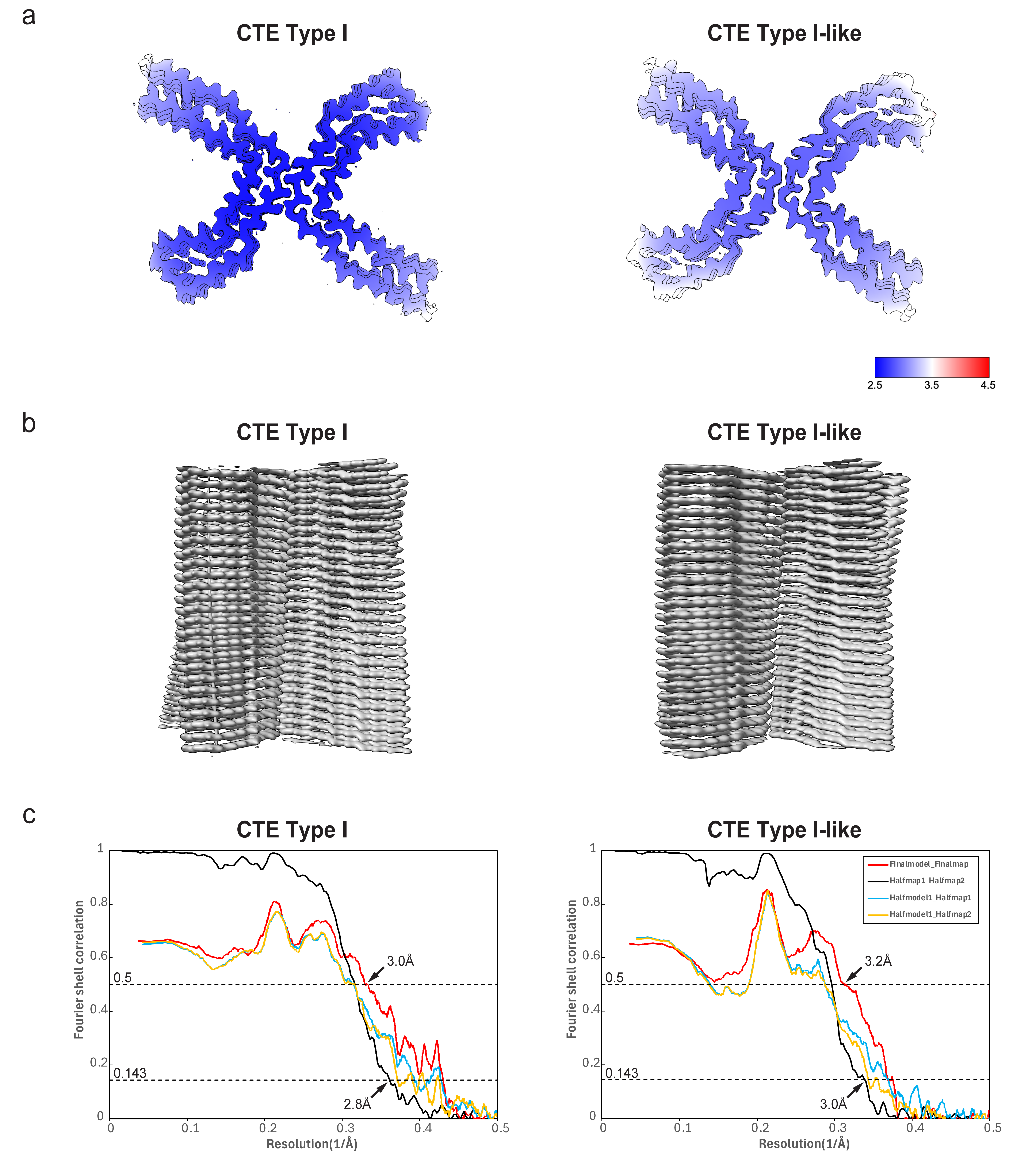

**Fig. S1** Quality assessment for the CTE Type I and CTE Type I-like reconstructions from EMPIAR-10313

(a) Central cross-sections of the post-processed maps colored by local resolution (color scale, 2.5–4.5 Å). (b) Side views of the final refined maps rendered along the helical axis. (c) Fourier shell correlation (FSC) curves showing half-map FSC (black) and model–map FSC (red, final model versus final map; blue and orange, model refined against half-map 1 compared with half-map 1 or half-map 2). Nominal map and model resolutions are indicated at FSC = 0.143 and 0.5, respectively.

### Table S1 Cryo-EM reconstruction, model refinement, and validation statistics

| **Data collection and processing** | | |  | **Model Refinement** | **CTE Type I** | **CTE Type I-like** |
| --- | --- | --- | --- | --- | --- | --- |
| Data source / Accession | EMPIAR-10313 | |  | Model resolution (Å, FSC=0.5) | 3.0 | 3.2 |
| Original report | Falcon et al., Nature 568:420–423 (2019) | |  | Map sharpening B factor (Å²) | -59.72 | -71.82 |
| Obtained images | 5,303 aligned micrographs | |  | Model composition |  |  |
| Obtained coordinates | TypeI_particles.star | |  | Non-hydrogen atoms | 7952 | 7952 |
| Physical pixel size (Å) | 1.055 | |  | Protein residues | 1036 | 1036 |
| Re-extracted particles (no.) | 198,298 | |  | Waters | 0 | 0 |
| **Helical reconstruction** | **CTE Type I** | **CTE Type I-like** |  | B factors (Å²) |  |  |
| Symmetry imposed | C1 | C1 |  | Protein | 99.65 | 50.73 |
| Final particles (no.) | 33,415 | 42,256 |  | Waters | - | - |
| Map resolution (Å, FSC=0.143) | 2.78 | 2.96 |  | R.m.s. deviations |  |  |
| Helical twist (°) | 179.4 | 179.4 |  | Bond lengths (Å) | 0.002 | 0.005 |
| Helical rise (Å) | 2.37 | 2.37 |  | Bond angles (°) | 0.464 | 0.562 |
| EMDB | 78521 | 78522 |  | Validation |  |  |
|  |  | |  | MolProbity score | 2.21 | 2.20 |
|  |  | |  | Clashscore | 7.64 | 8.50 |
|  |  | |  | Ramachandran plot |  |  |
|  |  | |  | Favored (%) | 95.83 | 97.22 |
|  |  | |  | Allowed (%) | 4.17 | 2.78 |
|  |  | |  | Outliers (%) | 0 | 0 |
|  |  | |  | PDB | 37VX | 37VY |
